# PhosPiR: An automated phospho-proteomic pipeline in R

**DOI:** 10.1101/2021.09.14.460225

**Authors:** Ye Hong, Dani Flinkman, Tomi Suomi, Sami Pietilä, Peter James, Eleanor Coffey, Laura L. Elo

## Abstract

Large-scale phospho-proteome profiling using mass spectrometry (MS) provides functional insight that is crucial for disease biology and drug discovery. However, extracting biological understanding from this data is an arduous task requiring multiple analysis platforms that are not adapted for automated high-dimensional data analysis. Here, we introduce an integrated pipeline that combines several R packages to extract high-level biological understanding from largescale phosphoproteomic data by seamless integration with existing databases and knowledge resources. In a single run, PhosPiR provides data clean-up, fast data overview, multiple statistical testing, differential expression analysis, phospho-site annotation and translation across species, multi-level enrichment analyses, proteome-wide kinase activity and substrate mapping and network hub analysis. Data output includes graphical formats such as heatmap, box-, volcano- and circos-plots. This resource is designed to assist proteome-wide data mining of pathophysiological mechanism without a need for programming knowledge.

## INTRODUCTION

Protein phosphorylation is a reversible post-translational modification (PTM) catalyzed by protein kinases of the transferase class [1]. Since its discovery in 1932, numerous studies have highlighted the importance of phosphorylation as a central regulatory process in cells [2]. High numbers of often tightly interconnected phosphoproteins participate in cell signaling and all aspects of cellular function from proliferation and differentiation to metabolism and neurotransmission, to name a few [3] [4] [5]. Phosphorylation is the most abundant “signaling” post-translational modification exceeding ubiquitination, methylation, and acetylation [6]. To understand how complex phosphorylation changes, especially shifts introduced by pathophysiological states coordinate function, systems-level phosphoproteomics study becomes necessary [6]. Advanced mass spectrometry methods enable high throughput measurement of phosphoproteomes [7], however traditional downstream analysis does little beyond phosphopeptide identification and quantification. Recent developments in R packages have taken advantage of protein phosphorylation databases and annotation advances, thereby supporting the creation of an analysis tool that can better exploit phosphopeptide data.

Here we introduce PhosPiR, a pipeline which takes advantage of available open-source tools for a complete downstream analysis of mass spectrometry-derived data after phosphopeptide identification. No programing knowledge is required to run the pipeline. Our workflow consists of peptide quality control, data overviewing with histogram, boxplot, heatmap, and PCAs, data annotation utilizing UniProt and Ensembl database, differential expression analysis including four statistical test options and post-hoc testing, phospho-site translation across species, four enrichment analyses for phosphoproteins, PTM-SEA (post translational modification set enrichment analysis) for phosphopeptide, kinase analysis, network analysis and hub analysis (Figure 1). The pipeline is accompanied by video tutorials and we exemplify the functionality of the tool using previously published largescale phosphoproteomic study of circadian clock changes in synaptoneuroscomes [8].

**Figure 1.**
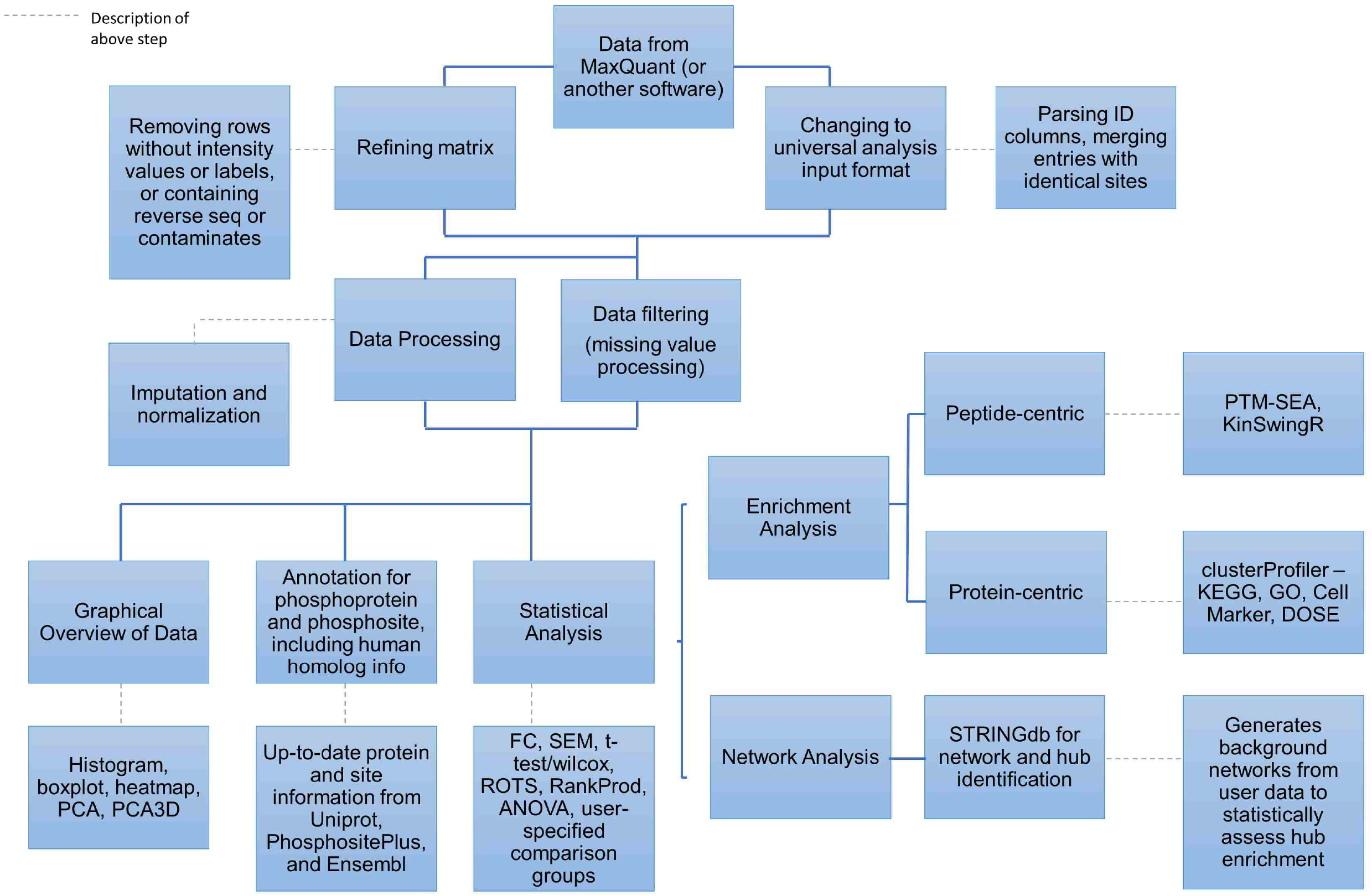
Flowchart overview of pipeline architecture. The main pipeline steps are outlined. Information on software packages utilized and/or information on the approach used is provided in adjoining boxes.

## METHODS

### Input data

PhosPiR accepts output files directly from MaxQuant or preprocessed files from similar MS data processing platforms that provide peptide sequence identification and intensity data on post-translational modifications (PTMs) [9]. The input file for PhosPiR is “Phospho (STY)Sites.txt” from the “combined/txt” folder within MaxQuant. In case another spectra analysis tool is preferred instead of MaxQuant, such as Progenesis, Spectronaut, openMS, or PEAKS, PhosPiR has an “Other” option for input file. The following steps are all outlined in the user support video found at our GitHub page. The data input should follow the following format:

1. File format should be .csv.
2. The first 6 columns should contain information as follows. Column 1 contains the UniProt protein ID, column 2 contains the protein description, column 3 displays gene name, column 4 shows the phosphorylated amino acid residue, column 5 records the phosphorylation site position within the protein, and column 6 shows the sequence window. The sequence window should have the phosphorylation site in the center position and extend in both directions by at least 7 amino acids. The order of the columns is crucial and should not be randomized.
3. Column 7 onward should contain sample intensity values. Each column corresponds to 1 sample, and each row corresponds to 1 phosphorylation pattern. Intensity values should not be in log scale.
4. Missing values should be written as “NA” or 0. Intensity columns should be numeric columns.

An example input file is shown in Supplementary Table 1. PhosPiR also supports proteomics data with the “Other” option for non-phospho proteomics data. In this case, when following the above format, column 5 and 6 should be marked “NA” for every row. To start the workflow, simply run the “run.R” script in R program. R can be downloaded from https://www.r-project.org/.

### Filtering and normalization setup

Filtering and normalization steps are implemented as data quality control. It is important to understand the data and make educated choices here to bring forth the most reliable result. Filtering removes outlier rows and/or columns with excess missing values. The user can always choose not to filter. For normalization, there are 3 choices, normalize and impute, only normalize, or neither. proBatch package [10] and MSImpute package [11] are utilized for normalization and imputation, respectively. Median normalization is performed when “only normalize” is chosen. Quantile normalization and low-rank approximation imputation is performed when “normalize and impute” is chosen. These normalization methods were chosen as they have proven successful with phosphoproteomics [8, 10, 12]. Imputation requires at least 4 non-missing values per row, if the input data does not satisfy this requirement, the user will be forced to choose one of the other two options.

### Other information setup

Organism information, sample group information, and comparison information needed to be setup by the user. PhosPiR provides a prompt window accompanied by a guide for the setup process. For organism selection, the user should highlight an organism from the organism list, or if the organism is not available from the list, select “Other” from the list, then input the scientific name of the source organism. A few analysis steps are only available for human, mouse, or rat data.

For sample group information, the user will be asked how many group classifications are found in the data, for each classification, a brief description is entered e.g. treatment or genotype, followed by group order. The group order is entered by providing a single number for each sample that represents its group classification, separated by a comma. For example, if a dataset contains 10 samples, the first 5 from a patient group designated “1” and the other 5 from a healthy group designated “0”, the user would enter “1,1,1,1,1,0,0,0,0,0” in the prompt, and so on for additional groups. Multiple group comparisons can also be setup. In this case, the user chooses the group classification first, and then enters e.g. “1,2” to specify the groups that should be compared against each other. The process can be repeated for as many group comparisons are wished. Lastly, the user can select whether to run all analysis steps, or only annotation/overview figure step. After setup, PhosPiR will run the analyses automatically, however, between analyses, step-specific choices will be given to the user, and the pipeline will not continue until the user has responded.

### Overview Figures

Several figures will be plotted automatically in order to provide an overview of the data distribution. Boxplot and histograms are suitable visuals for comparing sample distributions. Heatmaps, PCA with k-means clustering, and 3D PCA plots will automatically display the results from unsupervised clustering of the data, providing informative biological patterns from the data. Log 2 intensity values are automatically generated and used for overview figures. A few packages are utilized for this step. Boxplot and histogram are plotted with “ggplot2” [13]; heatmap is plotted with “pheatmap” [14]; PCAs and 3D PCA are plotted with “rcdk” [15], “fingerprint” [16], “vegan” [17], “rgl” [18], “FactoMineR” [19], “factoextra” [20], “plot3D” [21], and “magick” [22].

### Data annotation

A data annotation step utilizes information from all organisms found in the Ensembl database to identify for example reviewed accession and phosphorylation site position, Entrez ID, genome position of the protein, human ortholog genome position, accession, identify score and sequence alignment, protein pathology, expression, posttranslational modifications, subcellular location, and links to publications containing information on the protein in question. For each unique protein i.d., this information is extracted from both the Ensembl and Uniprot database. For non-human organisms, the human ortholog information is also included for comparison. Due to the long run time, the user has the choice to opt out of UniProt and human ortholog information mining.

Non-human organism data usually has many unreviewed accessions within the dataset. Some databases such as PhosphoSitePlus, host site information based on Swiss-Prot accessions, and does not include unreviewed accessions. This results in difficulties matching the input phospho-site identity to the database’s information. PhosPiR solves this issue by identifying the Swiss-Prot accession for the protein and aligning the sequences to generate the equivalent reviewed phospho-site position. This reviewed site information can be used for database searches by data annotation tools, thereby maximizing the identification of associated biological information. Human ortholog information allows for direct comparison of model organism data to human information. Pairwise alignment to human ortholog protein sequences allows the user to identify orthologous phosphorylation sites in human for any site of interest identified in their organism. Sequence alignment should only serve as a reference. While it should be accurate for alignments with high identity scores, its practicality decreases as sequence identify score decreases. The following packages are utilized for this step: “biomaRt” [23], “Biostrings” [24], “GenomicAlighments” [25], “protr” [26], and “UniprotR” [27].

### Differential expression analysis

Statistical tests are performed based on the group comparison setup. The user selects whether or not the data is paired in the given comparison and is offered a choice of statistical tests. For two group comparisons, fold change is automatically calculated. The fold change direction is determined by the group number, where the group with the larger number is the numerator. Group numbering should therefore take this into account. Four statistical tests T-test, Wilcoxon signed-rank test, reproducibility-optimized test statistic (ROTS) and rank product test can be chosen. T-test should not be chosen for nonparametric data. All tests can be selected if desired. Each test will yield a p-value and an FDR value for each data row. For a multiple group comparison, ANOVA with post-hoc Tukey HSD Test will be performed if the groups are not paired, and linear mixed effect modeling (LME) is performed if groups are paired. Next, the user can set thresholds based on p-value or FDR for example. The significant lists will then be used as input for enrichment and network analyses. The user can choose volcano plots to visualize the statistical results. Multiple comparisons (maximum 4) can be plotted together. Based on the selected statistical cut-off, significant entries in the plot will be labeled if the number of significant changes is less than 60 in total. ROTS analysis utilizes the “ROTS” package [28], “RankProd” [29] performs the rank product analysis, “multcompView” [30], “lsmeans” [31], and “nlme” [32] are ultilized for LME modeling. “ggrepel” [33] and “gridExtra” [34] are utilized in addition to “ggplot2” for volcano plots.

### Enrichment Analysis

Enrichment analysis is performed on both phospho-protein intensity data and phospho-site data. Protein level enrichment utilizes the “clusterProfiler” package [35]. This powerful analysis tool enables gene ontology (GO) enrichment, Kyoto Encyclopedia of Genes and Genomes (KEGG) pathway enrichment and cell marker enrichment, and for human data, disease-association enrichment. For KEGG analysis, it automatically utilizes the current latest online version of KEGG database which for example includes the COVID-19 pathway from the newest release (v.98). All the listed enrichment analyses are performed for each significant list created earlier. Both universal background and dataset background are applied for separate analyses. Only enrichments with significant entries are recorded as a result. Phospho-site enrichment utilizes the “PTM-SEA” (PTM-signature enrichment analysis) tool and its library PTMsigDB [36]. PTMsigDB curates detailed post-translational modification (PTM) information based on perturbation-induced site-specific changes, such as the direction of phosphorylation change upon a signaling event, and the affected signature sets that are collectively regulated. PTM-SEA analysis is performed on the entire dataset for all group comparisons. It is available for human, mouse, and rat.

### Kinase Substrate Analysis

A kinase–substrate relationship analysis is performed with the “KinSwingR” package [37], on each group comparison. It predicts a kinase-substrate interactome based on a library of motifs, and then integrates the fold change direction and P-value from the statistical results, to calculate a normalized swing score. The score resembles a z-score which predicts kinases’ activity changes. The P-value is calculated to determine the significance level based on a permutation test. PhosPiR utilizes the latest kinase information from the PhosphoSitePlus human, mouse, and rat data for customized kinase library [38] instead of the kinase dataset included in KinSwingR, which has become outdated and is only available for human.

A kinase-substrate network Circos plot is automatically created with the “circlize” package [39] which shows the top 250 significant substrates. Kinases are connected with edges to their specific substrate sites with phosphorylation sites grouped by phosphoprotein.

### Phosphoprotein/Protein Network Analysis

The phosphoprotein network is built using the “STRINGdb” package [40]. The STRING tool uses its own protein:protein interaction database which is used in PhosPiR to build an interaction network from each significant data list. Only interactions with greater than 0.4 confidence score (ranging from 0-1) are included in the STRING network figure (e.g. Supplementary Figure 1). The user can choose to identify hubs from within each network, and a hub interaction-enrichment score is also calculated. Hub phosphoproteins represent highly connected proteins within the network and are therefore likely to be functionally informative. The user can also choose from two cut-offs for defining hubs: the top 10% highest interactions, or an interaction count of one standard deviation above the mean. The hub interaction enrichment is calculated by generating 1000 control networks for each hub and comparing hub interactions in control networks to the query network. Both p-value and FDR are presented in the result.

## RESULTS

### Description of example setup and output

To demonstrate the functionality of PhosPiR, we analyzed synaptoneurosome phosphoproteome data from Brüning et al., 2019 [8]. In the original study, the authors studied the phosphorylation changes over time in synaptic terminals (otherwise known as synaptoneurosomes) from sleep-deprived mice and control mice. Here, we compared the overall differences in synaptoneurosome protein phosphorylation from mice while awake or asleep under control or sleep-deprived conditions.

The original study processed the synaptoneurosome phosphopeptide data with the EasyPhos platform. The sleep wake cycles were controlled as follows: mice were kept in a light:dark 12 hours cycle, synaptoneurosomes were taken every 4 hours (n=4) in a single day for sleep-deprived mice and baseline mice, totaling to 48 samples [8]. For the re-analysis of this data, we downloaded the raw MS files from the PRIDE database with identifier PXD010697. Taking the original data preparation as a reference [8], we preprocessed the raw data in a similar fashion using MaxQuant 1.6.17.0 [9] and Perseus 1.6.7.0 [41] (Supplementary Table 1). When inputting the preprocessed data into PhosPiR, “Neither” was selected for normalization and imputation, as it is done outside of the pipeline.

The folder organization of the output files from PhosPiR is shown schematically in Supplementary Figure 2. The *Group Information* folder describes the group comparisons as setup by the user. Examples of group information can be found in Supplementary Tables 2 and 3, respectively. In the Overview Figure folder, the overall data distribution is visualized in several ways including histograms, heatmaps, PCA, and boxplots plots. An example is shown in Supplementary Figure 3. The *Statistical Analysis* folder presents the statistical data including significance threshold for several tests as well as volcano plot output of significantly changing phosphopeptides or proteins, for each comparison and for every statistical test selected. Examples of statistical analysis output are shown in Figure 2A and Supplementary Table 4. The *Enrichment* folder stores the protein-centric result on cell marker enrichment, GO enrichment and KEGG enrichment (Figure 2B and Supplementary Table 5), and peptide-centric PTM-SEA enrichment result. PTM-SEA data is stored in the *PhosphoSite enrichment* subfolder, which includes converted .GCT input files and PTM-SEA analysis results, one per comparison, each in its own folder. Example rank plots of phosphorylation signatures can be found in Figure 2C. *Kinase Analysis* folder includes predicted significant kinases analyzed from KinSwingR, and resembled by swing scores, similar to z-scores for kinase activity change, and p-values, which indicate the significance of this change. Kinase activities are evaluated from the entire set rather than from the significant list. Examples of motif diagrams and kinase swing score output can be found in Figure 3 and Supplementary Table 6, respectively. The *Network* folder provides interaction figures of kinases to substrates spinning off from kinase analysis (Figure 4), also provides output from the STRING database network analysis and hub significance analysis. Examples are shown in Supplementary Figure 1 and Figure 5. The *Annotation* folder includes important ID information, UniProt database information, human homolog information and sequence alignment for all proteins as well as phosphorylation sites from the dataset. An example of human homolog ID information and UniProt database information can be found in Supplementary Table 7 and 8, respectively.

**Figure 2.**
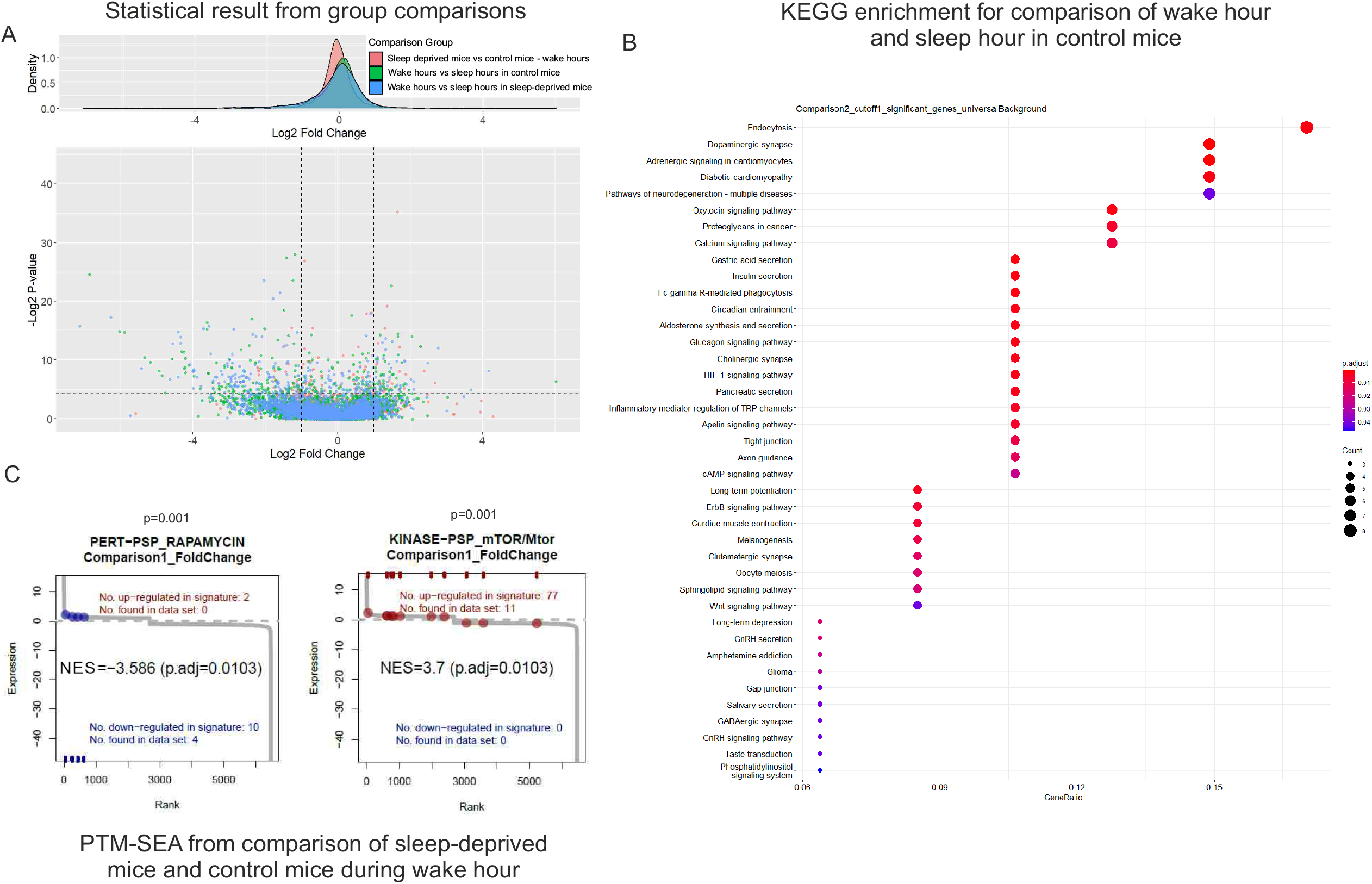
Sample output from the *statistical* and *enrichment* analysis feature of PhosPiR. Phosphoproteome changes in synaptoneurosomes of sleep-deprived verses normal sleep cycle mice. **A**. Normalized phosphoproteomic data was loaded to PhosPiR. Automated *statistical analysis* was done on user-defined group comparisons for up to 4 statistical tests and visualized as volcano plots and csv files. A representative volcano plot is shown for the rank product statistical analysis output. Significant proteins are labeled in the volcano plots only when there are ≤ 60 significant datapoints, otherwise the labels overlap. In this example the number of significant hits are > 60. Every volcano plot is accompanied by a csv file providing detailed numerical output from all statistical tests, including UniProt and gene accession identifiers. **B**. PhosPiR performs several enrichment analyses on the data e.g. GO, cell marker and KEGG enrichment analyses. The *KEGG analysis* output is shown from the comparison between wake and sleep time synaptoneurosomes phosphopoteome, as an example. **C**. Phosphosite enrichment using the *post-translational modification set enrichment analysis (PTM-SEA)* compares synaptoneurosomes from sleep-deprived mice and control mice during wake hours. Enrichment P-values and FDR (adjusted p-value) are indicated. This analysis highlights synaptic upregulation of mTOR pathway phosphorylation in sleep-deprived mice. Information on specific proteins and regulated sites are found in the accompanying csv file in the Enrichment\PhosphoSite enrichment folder.

**Figure 3.**
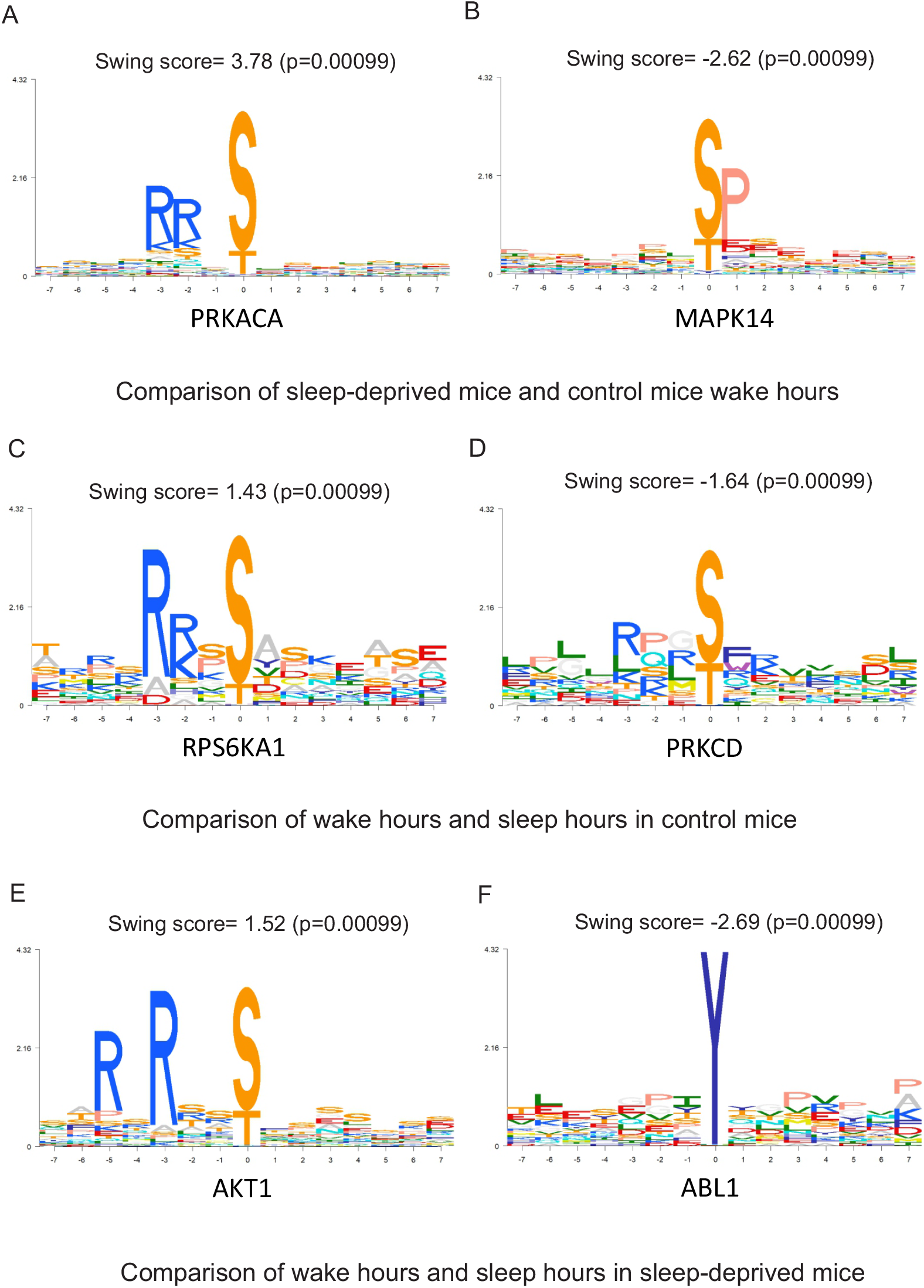
PhosPiR utilizes the KinSwingR tool to predict increases or decreases in kinase activities for defined group comparisons. PhosPiR integrates the KinSwingR tool which assesses local connectivity (swing) of kinase-substrate networks. Automated output from PhosphoPiR *kinase analysis* predicts regulated kinase activity based on identified substrate motifs. The final swing score is a normalized and weighted score of predicted kinase activity. Swing scores, positive and negative represent the direction of kinase activity change. Representative output tiffs are shown and accompanying csv file (ComparisonX_swingscore) is found in the *Kinase analysis* output folder.

**Figure 4.**
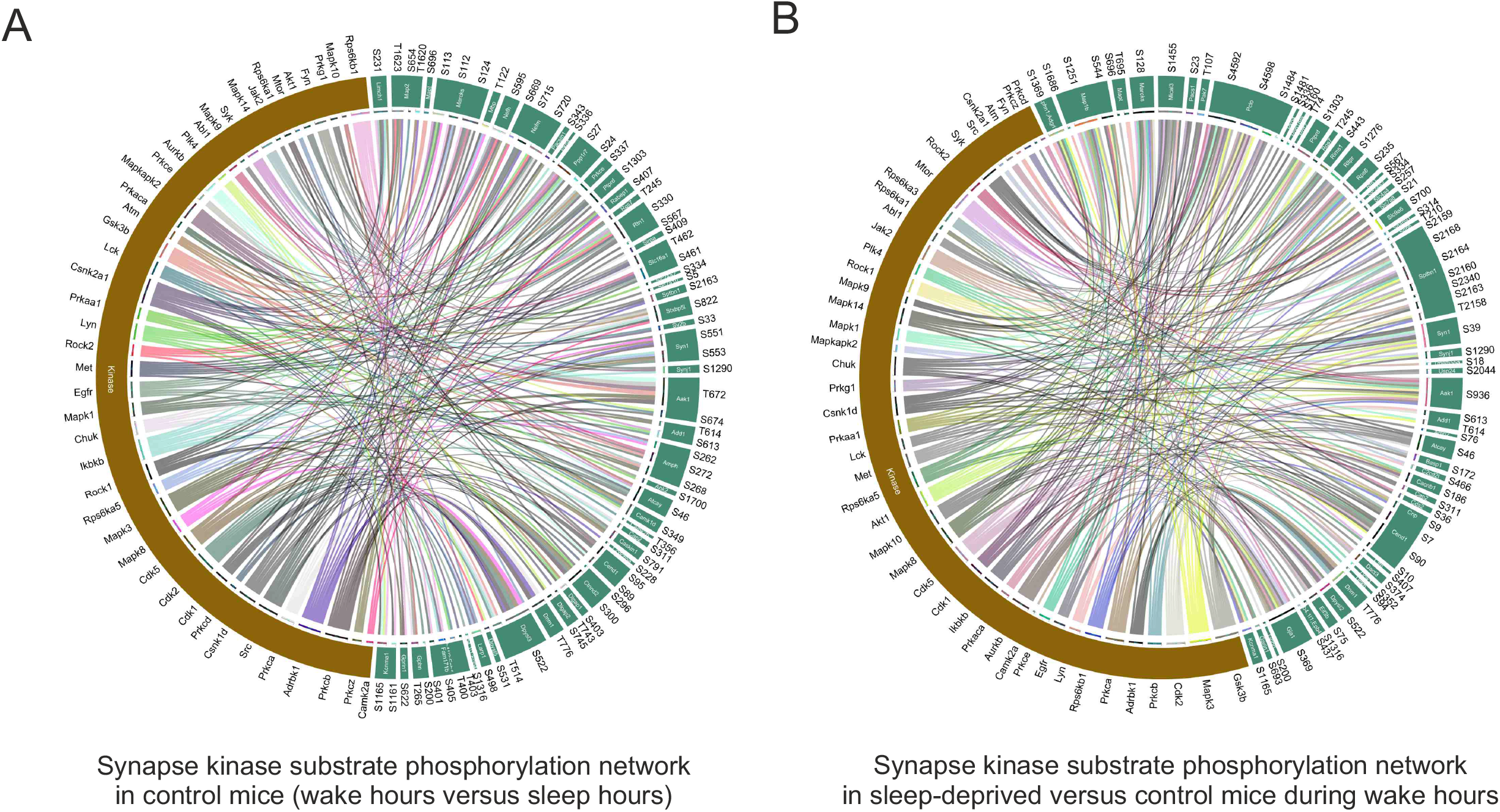
Kinase substrate network prediction tool. PhosPiR performs a proteome-wide *kinase analysis* using the KinSwingR package as shown (Fig. 3). The PhosPiR *Network Analysis* tool finds the top kinase-substrate relations and presents them in a Circos plot. **A, B**. Predicted kinase-substrate connections from the significantly changing data for group comparisons (**A**) wake hours verses sleep hours from control mice and (**B**) sleep deprived vs control mice during wake hours are shown. Colored ribbons link the kinase of interest with the substrate phosphorylation site that is significantly changed in the comparison. Predictions rely on known kinase-substrate phosphorylation sites. Only the top 250 most significant kinase-substrate relationships are plotted, to facilitate labeling. All output data is available in the accompanying csv file ComparisonX_significant_kinaseNetwork.

**Figure 5.**
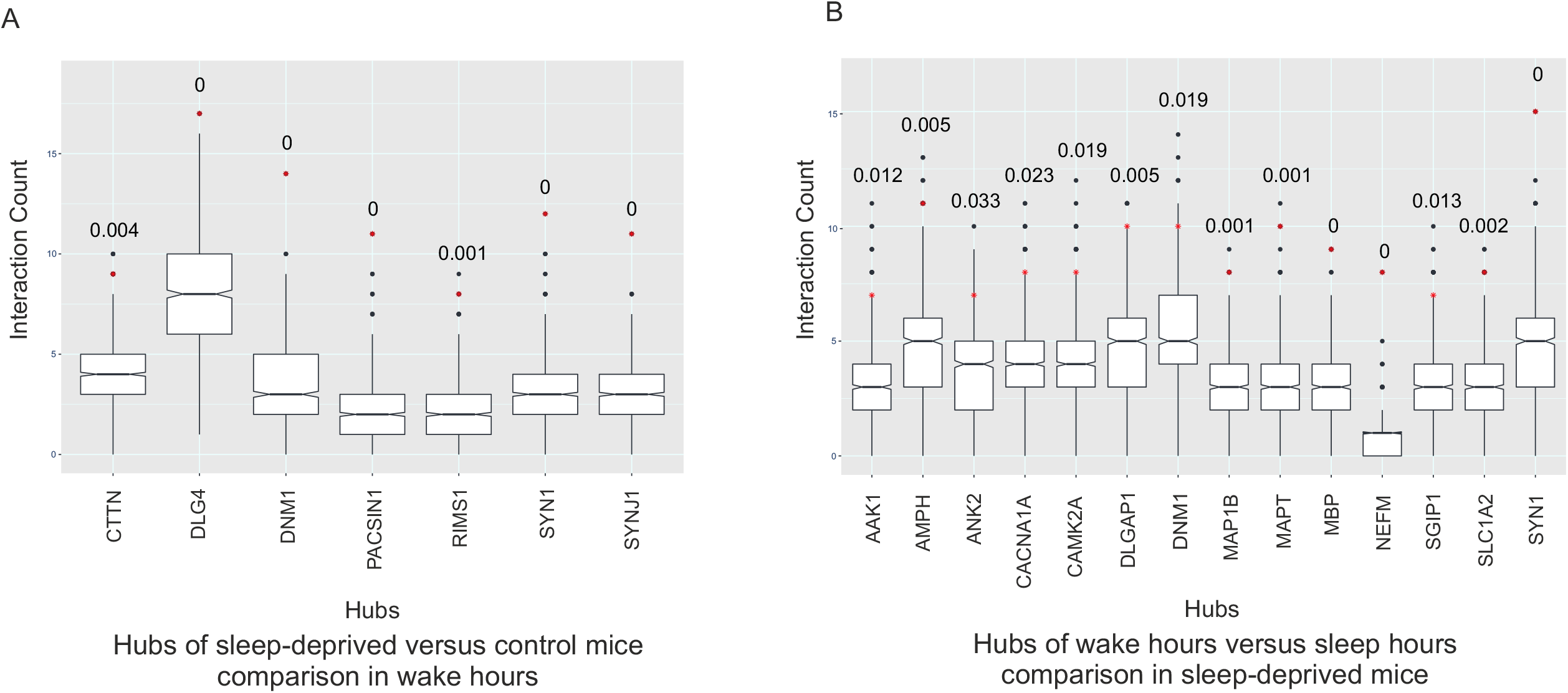
PhosPiR identifies network hubs based on protein:protein interactions. Sample output from the *Network Analysis* tool hub analysis is shown in **A** and **B**. Hubs are defined as proteins with interaction number > 1 standard deviation from the mean. Hub significance is calculated from the number of interactions within the data set compared to 1000 equal sized background datasets randomly generated from the total data. The hub interaction count in the background dataset is shown as a boxplot, and interaction count (hubness) in the *target* network is indicated by a red star. P-values calculated from the permutation test are indicated above the boxplots. Hubs from comparisons of sleep deprived vs control mice during wake hours, and wake hours verses sleep hours from sleep-deprived mice are shown in **A** and **B**, respectively.

### Description of example results

The MS data used here [8], incorporated intensity data for 13,634 phosphosites from which, 8,386 remained after filtering. Among these, PhosPiR identified 61 known disease associated phosphorylation sites, and 256 known regulatory sites using the automatic detection annotation tools (Supplementary Tables 9 and 10). The group comparisons (control verses sleep-deprived, and wake period verses sleep period with or without sleep-deprivation), identified 367 significantly changing phosphorylation sites with fold change ≥ 2, and Rank product FDR of <0.05. These results can be seen from volcano plot and csv file output (Figure 2A, Supplementary Table 11). Interestingly, the proteins with significantly altered phosphorylation between wake and sleep time were enriched for changes on the dopaminergic synapse pathway, as shown in Figure 2B. Thus, significant phosphopeptide changes were identified for voltage gated ion channels VGCC, VSSC and Cav2.1/2.2, and for signaling proteins PLC, PKC and CamKII (Figure 2).

In the phosphosite-centric enrichment analysis, the signature set “rapamycin” was 40% downregulated and the “mTOR” signature set was 14% upregulated, in sleep-deprived synaptoneurosomes (Figure 2C), consistent with known negative regulation of mTOR by Rapamycin [42]. This demonstrates the utility of the PhosPiR pipeline to make functionally accurate predictions as the mTOR pathway is known to regulate sleep-deprivation induced responses [43, 44]. Moreover, as the PhosPiR identifies specific phosphorylation sites from these signature sets, as well as their regulatory function, where documented (Supplementary Table 12), far more detailed mechanistic insight can be gained using PhosPiR’s integrated approach than would be possible with a stand-alone phospho-site analysis. This is further supported by the PhosPiR kinase activity analysis, which uses the KinSwingR package to predict kinase activity changes (Figure 3, Supplementary Table 13).

Examples of identified kinase substrates that undergo altered phosphosphorylation at synapses during wake verses sleep hours, or following sleep-deprivation, are shown in the Circos plots in Figure 4A and B. In Figure 4A, neurofilament M (NEFM) which shows the greatest fold change in phosphorylation (−115x) can be seen to be regulated by SRC, ADRBK1 (GRK2), CSNK1D and PRKCD in synaptoneurosomes when comparing wake hours to sleep hours. Examination of the corresponding statistics file reveals that RPS6KA1 has the most increased activity among kinases based on motif phosphorylation, and PRKCZ shows the largest decrease in activity during wake hours in sleep-deprived mice based on observed phosphorylation changes on known kinase motifs (Figure 4B). This figure also highlights mTOR, this time as one of the kinases with the largest number of substrates that undergo altered phosphorylation in synaptoneurosomes from sleep-deprived mice. The precise substrates; adhesion G-potein coupled receptor L1 (LPHN1), cell cycle exit and neural differentiation protein 1 (CEND1), piccolo (PCLO), F-actin mono-oxygenase (MICAL3), spectrin beta chain (SPTBN1), MARCKS, GJA1 and MAP1B, and their altered phosphorylation sites, can be read directly from the plot (Figure 4B).

The hub analysis of protein phosphorylation in synaptoneurosomes during wake time verses sleep time has identified that the NMDA receptor subunit GRIN2B was a highly significant signaling hub (Figure 5A). This is consistent with several reports pointing to NMDAR in sleep regulation especially in the context of autoimmune encephalitis-induced sleep disturbance [45, 46]. Similarly, SHANK3 was highly connected to the changing phosphoproteins, consistent with its reported action in the control of circadian rhythm [47]. Synapsin I (SYN1), a neurotransmitter release regulatory protein, was also highly networked with the wake cycle phospho-proteins. SYN1 has previously been associated with synaptic changes following sleep deprivation [48]. Conversely, in sleep-deprived mice, there were fewer hubs overall consistent with the finding of Bruning et al, which showed that phosphorylation cycling was reduced upon sleep deprivation [8]. Nonetheless, synapsin I, neurofilament (NEFM) and MAPT showed increased connectivity to the regulated phosphoproteins (Figure 5B). MAPT phosphorylation has been shown to increase upon sleep deprivation stress [49]. Thus, PhosPiR automated analysis identified known regulators of sleep/wake cycle and sleep deprivation stress in synaptic terminal preparations from mouse brain. Moreover, PhosPiR provides site-specific and network information that can assist detailed parsing of mechanism.

As evident from the results, in addition to kinase-substrate prediction, kinase activity directional changes are also predicted by PhosPiR. Significantly, phosphorylated substrates are linked to kinases that are predicted to change activity, and the enrichment is checked with PTM-SEA. Moreover, as all identified sites are matched to their human homolog with pairwise alignment (Supplementary Table 14), analysis of pathological implications could be further confirmed from database search of aligned homolog sites. These are some of the highlights of PhosPiR.

## Discussion

PhosPiR is an automated pipeline that does not require any coding knowledge from its user. It integrates several new phosphoproteomic analysis tools such as PTM-SEA and KinSwingR into a single pipeline while it simultaneously translates phosphoproteomic data from model organisms to human in order to exploit a range of customized databases that facilitate identification of functionally relevant information.

Although all analysis steps are automatic, the pipeline retains flexibility through its setup options. The user is free to fully customize group comparisons. For example, in addition to the examples shown here, a time series analysis can also be included, by choosing the pairwise multi-group comparison option. All user options are presented with textboxes created using the “svDialogs” package [50] creating a straightforward and seamless experience for the user. Although the pipeline can be applied also to non-phospho proteomic data, several of the functions are specific to phospho-proteomics. For example, the ortholog alignment function, PTM-SEA enrichment analysis, kinase substrate analysis and the kinase network figure generation all rely on phosphorylation-site information. Among the unique features is the ortholog alignment tool that provides a human reference site for every single phosphorylation-site from mouse or rat. An important benefit of PhosPiR is that it integrates several packages, such as PTM-SEA that utilize customized, curated libraries that contain up to date information. The integrated “kinase analysis” function in PhosPiR not only predicts which kinases are linked to the input data, but also predicts activity changes based on the generated statistics. The “kinase network” function clearly labels substrate phospho-site position and organizes them by protein, thereby providing a clear visual summary for significant kinase substrate changes.

Many important functions of the pipeline come from recently developed, powerful R-packages, which the pipeline unifies and provides important customizations. For example, most of the included analysis packages require strict input formats and generate diverse output formats. PhosPiR utilizes packages such as “reshape2” [51], “vroom” [52], “openxlsx” [53], “textreadr” [54], “plyr” [55] and “cmapR” [56] to transform data between analysis steps, so that input requirements are satisfied, and output results are unified. Output figures are expanded from the originals, modified with packages such as “gplots” [57], “gridExtra” [34], and “RColorBrewer” [58] to be informative at a glance. Many more packages are utilized and listed within Method section, we wanted to include all of them to offer their functionalities and customizations to a wider audience of non-bioinformaticians. PhosPiR also supports a wide range of organisms. With the exclusion of the disease ontology semantic enrichment analysis (DOSE), kinase analysis and PTM-SEA which search only human data, whereas all other analysis steps support up to 18 organisms.

Our pipeline offers unique functionalities compared to even the most recent analysis packages such as PhosR [59], which provides a kinase analyses toolkit for bioinformaticians working in R coding language. In contrast, the PhosPiR workflow doesn’t require coding knowledge and can be performed by non-bioinformaticians. Furthermore, PhosPiR provides automated protein and site annotation from UniProt, Ensembl, and PhosphoSitePlus. The annotation files provide information on functionality and associated pathologies. Scientific references for identified functions are included in the output. Another unique feature of the pipeline is the protein-centric network and hub analysis, which provides aligned sequence information on human homolog when for example the input data is from a model organism. Finally, the annotations accompanying the kinase enrichment function (using the PTM-SEA database) takes into account directionality of phosphorylation change when identifying pathology and regulatory signatures. These many exclusive features enable users to study their data from multiple angles and distinguishes PhosPiR from existing phospho-proteomic data analysis software.

The current pipeline marks version 1.0. New functionalities are already in development. We aim to provide an automated result report in the next version and provide direct input support for results generated from more spectra analysis tools. The most time-consuming step of comparison setup will also be further optimized. We hope PhosPiR provides an opportunity for users with limited programming knowledge to experience great R packages for comprehensive functional prediction analysis with statistical support, from their phosphoproteomics data.

## Data availability statement

Example data is part of the study F. Brüning, S.B. Noya, T. Bange, S. Koutsouli, J.D. Rudolph, S.K. Tyagarajan, J. Cox, M. Mann, S.A. Brown and M.S. Robles, “Sleep-wake cycles drive daily dynamics of synaptic phosphorylation,” Science, vol. 366, pp. eaav3617, 10/11. 2019. It is available on PRIDE database, at https://www.ebi.ac.uk/pride/archive/projects/PXD010697.

## Software availability statement

The PhosPiR software will be available at the Github repository at the following site: https://github.com/TCB-yehong/PhosPiR. A short tutorial video can be accessed via link: https://youtu.be/8qvEStg28dQ.

